# Pre-search attentional focus supports learned suppression in visual search

**DOI:** 10.1101/2025.05.29.656755

**Authors:** Yinfei Zhou, Lishuang Wang, Xinyu Li, Jan Theeuwes, Benchi Wang

## Abstract

Salient distractors often capture our attention, disrupting ongoing tasks. Recent studies suggest that, through statistical learning, prior experiences regarding distractor locations can reduce distraction by suppressing their corresponding locations. However, the proactive neural mechanisms supporting this learned suppression remain unclear. Our findings demonstrate that participants learn to suppress locations that are more likely to contain distractors relative to other locations. Using frequency tagging in electroencephalography (EEG) recordings, we observed significantly different tagging responses between high- and low-probability locations, along with a general decrease in alpha power (8–12 Hz) prior to search onset. Notably, the higher tagging frequency power at high-probability locations suggests that participants allocated greater attentional focus to these locations in anticipation of the search. These results suggest that anticipatory attentional deployment precedes the suppression of high-probability distractor locations after the onset of visual search.

## Introduction

In everyday life, we embrace lots of visual information from the environment, but cannot process all of them at once. To act and behave in a goal-directed manner, we focus our limited resources on relevant information and filter out distracting information (Johnston & Dark, 1986). Although selective attention is an effective mechanism that determines what we see and act upon, our attention may be captured by salient distractors. One efficient way to suppress these distractors is learning from past experiences (regularities) regarding distractors (Theeuwes et al., 2022; Wang & Theeuwes, 2018a, 2018b) via a mechanism called statistical learning (SL, Turk-Browne et al., 2005). For example, behavioral studies have established that learning to expect where distractors are most probable during visual search can help suppress these distractors and facilitate upcoming search (Ferrante et al., 2018; Goschy et al., 2014; Leber et al., 2016; Sauter et al., 2018; Wang & Theeuwes, 2018a, 2018b, 2018c). While this effect has been established many times, the neural mechanism regarding learned suppression remains less clear, especially whether this type of suppression can be implemented *proactively*, by modulating different distractor locations separately.

To this aim, we adopted electroencephalogram (EEG) recordings in humans when performing the additional singleton task (Theeuwes, 1992), in which participants were required to search for a unique shape (target) while ignoring a salient distractor. Notably, the salient distractor was presented more often in one specific location than in other three locations in the learning condition, introducing statistical learning which should result in the suppression of the frequent distractor location. To track the pre-stimulus attentional distribution due to learned suppression, a technique called frequency tagging (Ding et al., 2006; Malinowski et al., 2007; Morgan et al., 1996; Muller et al., 2003; Zhigalov et al., 2019) was used to separately assess frequency tagging response to different distractor locations. If learned suppression operates through proactive attention mechanism, we expect corresponding neural activity would emerge before the stimulus onset. We also measured alpha activity simultaneously, as alpha-band activity is considered to be a neural signature of functional inhibition (Foxe & Snyder, 2011; Jensen & Mazaheri, 2010; Klimesch et al., 2006), and is strongly modulated by spatial attention (Sauseng et al., 2005).

## Methods

### Participants

Twenty-eight undergraduate students (20 women and 8 men with a mean age of 20 years, pre-determined based on Wang et al., 2019) participated in this experiment. The study was approved by the ethics committee at South China Normal University (2020-3-013), and all participants provided written informed consent before taking part. They were all right-handed, had normal or corrected-to-normal vision, and were financially compensated (¥70 per hour).

### Apparatus and Stimuli

Stimulus presentation and behavioral data collecting were controlled by custom scripts written in Python and run on a 27-inch computer with a 1920 × 1080 resolution and a refresh rate of 60 Hz. Participants were seated in a sound-attenuated and dimly lit laboratory at a viewing distance of 60 cm. All stimulus were presented against a black background (Red-green-blue [RGB]: 0, 0, 0) with a white fixation dot (RGB: 255, 255, 255, radius: 0.25°) presented in the center of the screen. Four gray (RGB: 128, 128, 128) placeholders (2° × 2°) were presented 4° away from the fixation. The search array consisted of four discrete stimuli with different shapes (one circle among three unfilled diamonds, or vice versa) superimposed on the gray placeholders. The circle had a radius of 1° and the diamond was subtended 2° × 2°. Each shape had a red (RGB: 255, 0, 0) or green (RGB: 0, 255, 0) outline, and a vertical or horizontal white line (0.2° × 1.4°) inside (see Fig. 1A).

**Figure 1.**
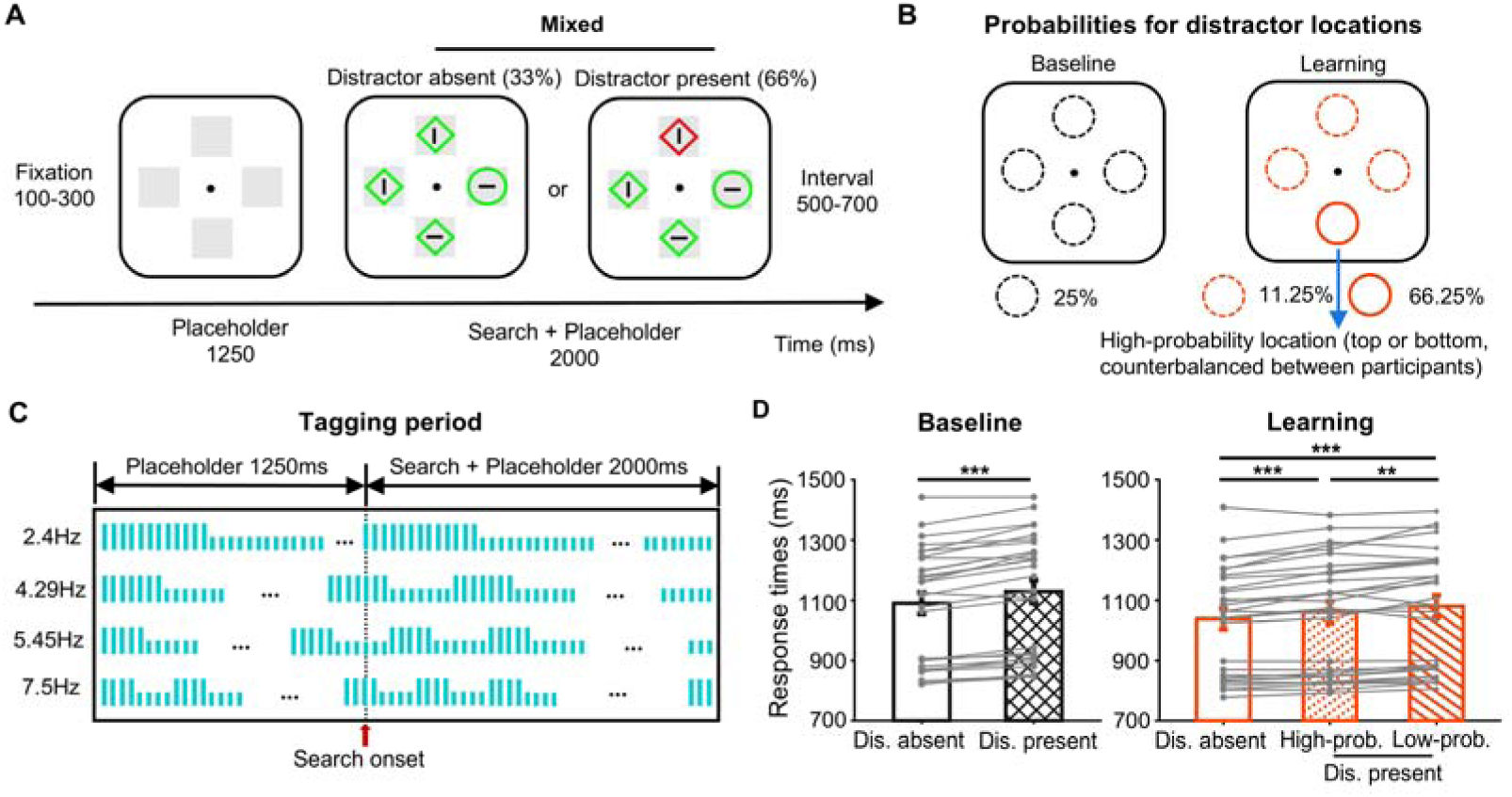
A) Experimental procedure. Each trial started with a fixation dot followed by four tagging placeholders for 1250 ms. Then the search array was presented with all tagging placeholders simultaneously for 2000 ms at different frequencies. Participants were asked to find the unique shape (in this case, the circle to the right) and report the orientation of the line inside (in this case, horizontal). Distractor present and absent trials were mixed within each block. B) Probabilities for distractor locations. In the baseline condition, the salient distractor was equally presented in four locations, with a probability of 25% each. In the learning condition, the salient distractor was presented in either the top or bottom location (which was counterbalanced between participants) with a probability of 66.25%, while was equally presented in other three locations with a probability of 11.25% each. C) Temporal sequence for different tagging placeholders at different frequencies, that is, 2.4 Hz, 4.29 Hz, 5.45 Hz, and 7.5 Hz (counterbalanced between participants). Each mini vertical bar represents a frame (16.7 ms), with the longer and short ones representing the onset and offset of the placeholders respectively. D) Mean response times (RTs) in the baseline and learning conditions. The gray solid dots represent individual RTs. Error bars show ±1 standard error of the mean (SEM). \*\**p* < .01, \*\*\**p* < .001.

### Procedure and Design

During the experiment, participants were instructed to search for a unique shape (a circle among diamonds or vice versa) while ignoring a salient distractor (red or green, which was noticeably different in color from the other search elements). Participants had to indicate the orientation (horizontal or vertical) of the line within the target shape. They were asked to respond as fast and as accurately as possible. Each trial began with a fixation dot (100-300 ms), followed by four gray placeholders that flickered (tagged) for 1250 ms. Subsequently, the search array was presented for 2000 ms, with those four gray placeholders continued to flicker. Notably, participants were explicitly instructed to ignore the tagging placeholders, ensuring their attention remained focused on the primary visual search task and not diverted to the tagging backgrounds. The inter-trial interval was between 500-700 ms (see Fig. 1A).

Before the experiment, participants received verbal instructions and completed 15 practice trials to be familiar with placeholder tagging and the experimental task. Each participant completed 5 blocks for the baseline condition and 10 blocks for the learning condition. Each block comprised of 120 trials, with 40 distractor-absent trials and 80 distractor-present trials. In the baseline condition, which was performed before the learning condition to avoid any contamination from learning, the salient distractor was equally presented in four locations, with a 25% probability each. In the learning condition, the salient distractor was presented in either the top or bottom location (which was counterbalanced between participants) with a probability of 66.25%, while was equally presented in other three locations with a probability of 11.25% each. The target was randomly selected from non-distractor locations in both conditions, resulting in no clue for participants to figure out where the target would be. More blocks for the learning condition were designed to examine whether any effect observed was due to that the baseline condition was tested firstly for all participants (see the last paragraph in the Results section for more details).

Notably, we deliberately chose not to present the high-probability location laterally because alpha lateralization effects could be confounded by bilateral electrode changes, rather than specifically reflecting modulation of the high-probability location. For instance, findings from Wang et al. (2019) suggest that any observed increase in alpha power contralateral to the high-probability location might partially arise from decreases in power contralateral to the low-probability location.

### Frequency Tagging

Frequency tagging refers to the presentation of visual stimuli that tag at specific frequencies, resulting in steady-state visual evoked fields which can be observed in EEG signals (Norcia et al., 2015). Here we employed four different frequencies (2.4 Hz, 4.29 Hz, 5.45 Hz, and 7.5 Hz) for each placeholder before and after the search display onset, allowing us to track attentional distribution over time. These frequencies were counterbalanced between participants. The criteria for selecting these frequencies were, 1) to avoid any overlap with alpha oscillation (8-12 Hz) which was one of the primary foci in our study; 2) to prevent any high frequencies (>12 Hz) that may cause discomfort for participants; 3) to ensure that each frequency was not a harmonic of another; 4) to ensure that the number of frames corresponding to the frequency is an integer. Our monitor’s refresh rate was 60 Hz, resulting in 25, 14, 11 and 8 frames for 2.4 Hz, 4.29 Hz, 5.45 Hz, and 7.5 Hz respectively (see Fig. 1B). These frequencies corresponding to different locations were counterbalanced between participants.

### EEG Recording and Preprocessing

EEG data were acquired using 64 Ag/AgCl active electrodes connected to BrainAmp amplifiers (Brain Products, Munich, Germany), placed according to the extended 10-20 system and digitalized at 500 Hz. Here electrodes T8 and FT10 were used to record the vertical electro-oculogram (EOG), and electrodes T7 and FT9 were used to record horizontal EOG; moreover, electrodes TP9 and TP10 were used to record signals from mastoids, and electrode Fz was used as on-line reference.

The data were re-referenced to the mean of left and right mastoids and high-pass filtered using a cut-off of 1.5 Hz (for independent component analysis [ICA] only) and 0.1 Hz (for final analyses). Continuous EEG was epoched from –2000 to 4750 ms relative to the placeholder onset. Malfunctioning electrodes were visually detected and temporally removed from the data; and a 110–140 Hz bandpass filter was used to capture muscle activity and allowed for variable z-score cutoffs per participant based on the within-subject variance of z scores. After trial rejection, ICA was performed on the clean electrodes only. Together with the vertical and horizontal EOG signals, we visually inspected and removed ICA components that captured eye blinks, eye movement, or other artifacts that were clearly not brain-driven activity. Afterwards, we interpolated the malfunctioning electrodes identified earlier. Offline preprocessing was performed using MATLAB v2016 (MathWorks Inc.), incorporating EEGLab2020_0 and custom MATLAB scripts.

### Time-Frequency Analysis

Preprocessed EEG data were broken into event-related epochs (–2000 to 4750 ms relative to placeholder onset; avoiding edge artifacts from wavelet convolution), and then were convolved with a set of Morlet wavelets with frequencies including 2.4 Hz, 4.29 Hz, 5.45 Hz, 7.5 Hz, 8 Hz, 9 Hz, 10 Hz, 11 Hz, 12 Hz, 20 Hz, 25 Hz, 30 Hz. The number of wavelet cycles was logarithmically spaced between 3 and 12 to have a good trade-off between temporal and frequency precision. This might result in a potential overlap between neighboring frequencies. However, the selected frequencies corresponding to different locations were counterbalanced between participants, preventing such potential overlapping effects. Because each frequency selected for the high-probability location was just one of several frequencies selected within the alpha band, making it unlikely that its activity alone can represent the full alpha-band activity. Furthermore, even when we varied the number of cycles (1.5, 1.6, 1.7, 1.8, 1.9, and 2 times the corresponding frequencies) to better differentiate between tagging frequencies and to keep the temporal resolution fixed, the results remained consistent (see Fig. S3 for details). The power within our focused time window (0 – 3250 ms) was then averaged across trials and decibel-transformed relative to a pre-stimulus period of −500 to −300 ms relative to the placeholder onset. We did not calculate phase-locked activity for further analysis because our task required participants to switch their attention across different search elements to search for the target. As a result, the neural activity produced by statistical learning during visual search was not phase-locked.

Based on previous studies, we selected PO3/4, PO7/8, and O1/2 for further analysis (Antonov et al., 2020; Wang et al., 2019). We were specifically interested in the tagging frequencies (2.4 Hz, 4.29 Hz, 5.45 Hz, 7.5 Hz), alpha frequencies (8 Hz, 9 Hz, 10 Hz, 11 Hz, 12 Hz), three control frequencies (20 Hz, 25 Hz, 30 Hz; which was far from the tagging frequencies and alpha oscillation, avoiding potential impact from these key frequencies), and their comparisons.

### Cluster-based permutation test

To correct multiple comparisons, we used a cluster-based permutation test against a null-distribution shuffled from 1000 iterations (following Monte Carlo randomization procedure). Specifically, one-sample t-tests were performed across participants for conditional differences against zero to identify above-chance activity, and time windows with t-values larger than a threshold (*p* = .05) were combined into contiguous clusters based on adjacency. The cluster statistics was defined as the sum of the t values within each cluster. The null distribution was formed by randomly permuting condition labels for 1000 times in order to get the largest clusters per iteration. Clusters were determined to be significant if the cluster statistics was larger than the 95^th^ percentile of the null distribution.

## Results

Participants performed a visual search task, in which they were instructed to search for a unique shape (target) while ignoring a salient distractor (see Fig. 1A). In the baseline condition, the salient distractor was equally presented at all four locations. In the learning condition, the salient distractor was presented more often in either the top or bottom location (which was counterbalanced between participants; high-probability location), while was equally presented in other three locations (low-probability location), producing learned suppression on salient distractors. Notably, four gray placeholders that tagged at different frequencies before and after the search onset, and participants were explicitly told to ignore the tagging background (see Methods for details).

### Behavioral results

Incorrect trials, as well as trials on which the response times (RTs) were faster than 200 ms and slower than 2000 ms, or no response was given within 2000 ms, were excluded from analyses (1.29%). Mean RTs for different conditions are presented in Fig. 1D. In the baseline condition, compared with distractor-absent trials (1090 ms), paired t-tests revealed that the mean RTs were significantly larger for distractor-present trials (1129 ms), *t*(27) = 7.55, *p* < .001, *d* = 1.43, indicating capture by salient distractors. The results on error rates mimicked those for RTs. The error rates were higher for distractor-present trials (5.1%) than distractor-absent trials (3.8%), *t*(27) = 3.92, *p* < .001, *d* = 0.74.

In the learning condition, with distractor condition (low-probability location, high-probability location and distractor-absent) as a factor, a repeated-measures ANOVA on mean RTs showed a main effect, *F*(2, 54) = 26.88, *p* < .001, partial η^2^ = 0.5. Subsequent planned comparisons revealed larger mean RTs for a distractor presented in the high-probability location (1060 ms), *t*(27) = 4.76, *p* < .001, *d* = 0.9, and the low-probability location (1079 ms), *t*(27) = 6.4, *p* < .001, *d* = 1.21, compared to the distractor-absent condition (1038 ms). Crucially, there was a reliable difference between the high- and low-probability locations, *t*(27) = 3.37, *p* = .002, *d* = 0.64, suggesting that salient distractor was suppressed after learning its high-probability location. No significant effect for error rates was observed, *F*(2, 54) = 1.22, *p* = .303, partial η^2^ = 0.04.

Overall, the results demonstrated the classic attentional capture effect by salient distractors (Lin et al., 2024; Theeuwes, 1992) and further indicated that this capture can be suppressed through the learning of statistical regularities regarding distractor locations (Wang & Theeuwes, 2018a, 2018b, 2018c). A recent study by Kerzel et al. (2022) suggested that reduced capture is not solely due to suppression of the high-probability location but also results from increased interference from distractors at low-probability locations (i.e., distractor rarity). In this study, larger capture effects were observed at low-probability locations compared to equal-probability locations, which correspond to the baseline condition in the present study. However, in our study, the capture effect was similar at low-probability locations (41 ms) and in the baseline condition (39 ms). This discrepancy may stem from differences in the experimental design. Unlike the between-subjects design used by Kerzel et al., (2022), our within-subjects design introduced the baseline condition before the learning condition, allowing participants to practice extensively before the learning condition. This practice likely reduced capture effects, particularly at low-probability locations. Nevertheless, this discrepancy does not undermine our conclusion that the high-probability location was suppressed, a finding consistent with Kerzel et al. (2022).

Notably, compared to previous studies that only four items had on display, we observed much longer RTs (e.g., Wang & Theeuwes, 2020). One possible explanation is that the tagging frames may have reduced the visibility of the shapes, making the task more challenging and contributing to the slower RTs observed.

### Frequency tagging and alpha oscillation

In Fig. 2A left panel, we depict the power spectra across frequencies averaged over the tagging period (from 0 to 3250 ms) for the baseline and learning conditions, as well as their differences. To validate the efficacy of frequency tagging and assess the role of alpha oscillation in the present study, we computed the average power for tagging frequencies (2.4 Hz, 4.29 Hz, 5.45 Hz, 7.5 Hz), alpha frequencies (8 Hz, 9 Hz, 10 Hz, 11 Hz, 12 Hz), and three control frequencies (20 Hz, 25 Hz, 30 Hz). Notably, we did not employ tagging for the search elements; instead, the placeholders (i.e., the stimulus background) were tagged continuously throughout the trial, with participants explicitly instructed to ignore them. The background (i.e., tagging placeholders) served as a distraction for the search task. Previous studies have shown that enhancing attention to a stimulus causes the suppression of unattended (distracting) stimulus simultaneously (Andersen & Muller, 2010; Müller & Hübner, 2002), thereby leading to reduction in frequency power when tagging the unattended stimulus. Thus, we hypothesized that the power for tagging frequencies would be lower than zero, as the tagging placeholders were disregarded throughout the trial. Consistent with this hypothesis, the modulation of the tagging frequencies (averaged across frequencies) was significantly lower than zero, *t*(27) = 2.64, *p* = .014, *d* = 0.5.

**Figure 2.**
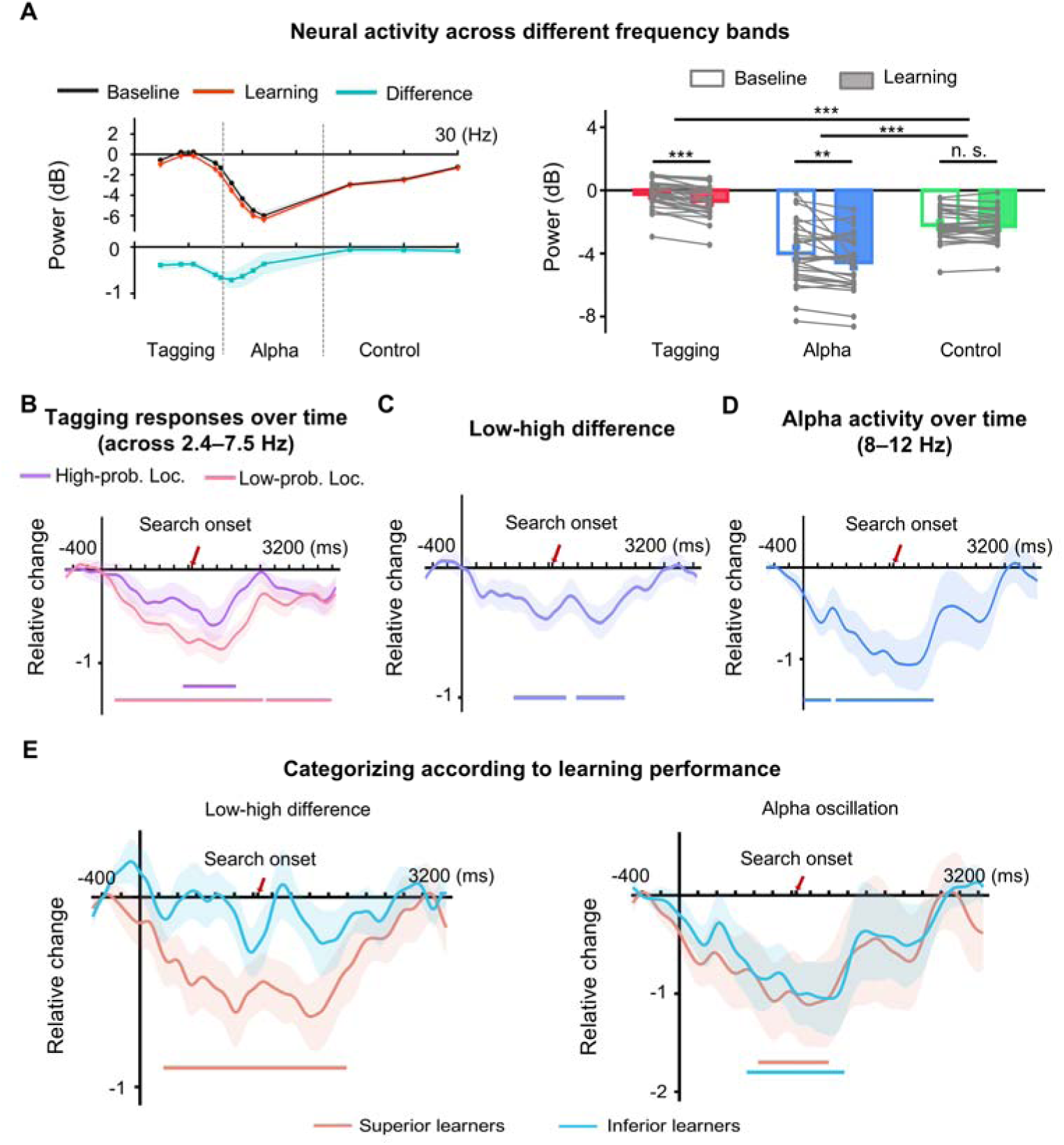
A) Left panel shows the power spectra across different frequencies averaged over tagging period (from 0 to 3250 ms) for the baseline and learning conditions, and their difference, including tagging frequencies (2.4 Hz, 4.29 Hz, 5.45 Hz, 7.5 Hz), alpha frequencies (8 Hz, 9 Hz, 10 Hz, 11 Hz, 12 Hz), and control frequencies (20 Hz, 25 Hz, 30 Hz). Right panel shows the averaged power for tagging-, alpha- and control frequencies in the baseline and learning conditions. The gray solid dots represent individual data. Error bars show ±1 SEM. \*\**p* < .01, \*\*\**p* < .001. B) Time-varied changes (i.e., power difference obtained by subtracting the power observed in the baseline condition from that in the learning condition) for tagging responses towards high- and low-probability distractor locations. C) The low-high difference (obtaining by subtracting tagging responses towards high-probability location from these towards low probability locations) indicate the ability to discern the high-probability location from other low-probability locations. D) Time-varied changes for alpha activity. E) The low-high difference (left panel) and alpha oscillation (right panel) for superior and inferior learners. Colored shadow areas represent ±1 SEM, and the horizontal lines represent significant clusters after cluster-based permutation test at *p* < .05.

Subsequently, with frequency type (tagging-, alpha- and control frequencies) and condition (baseline vs. learning) as factors, a repeated-measures ANOVA revealed significant main effects for frequency type, *F*(2, 54) = 90, *p* < .001, partial η^2^ = 0.77, and condition, *F*(1, 27) = 15.63, *p* < .001, partial η^2^ = 0.37, along with a noteworthy interaction, *F*(2, 54) = 6.11, *p* = .004, partial η^2^ = 0.19 (see Fig. 1A right panel). Further planned comparisons revealed that, in both baseline and learning conditions, suppression for tagging frequencies was weaker than that for control frequencies, both *p*s < .001, indicating tagging responses towards different locations following tagging placeholders. Additionally, alpha power exhibited a decrease relative to control frequencies, both *p*s < .001, signifying the involvement of alpha suppression in the present study.

Importantly, distinctions between baseline and learning conditions were evident for tagging frequencies, *t*(27) = 5.84, *p* < .001, *d* = 1.1, and alpha oscillation, *t*(27) = 2.99, *p* = .006, *d* = 0.57, but not for control frequencies, *t*(27) = 1.05, *p* = .302, *d* = 0.2. This shows that both frequency tagging responses and alpha oscillation decrease after the distractor location can be anticipated compared to the baseline condition.

### Tracking high- vs. low-probability locations over time

To explore the related neural activities associated with learned suppression, we initially calculated the power difference at each time point inside the tagging time window (0 – 3250 ms). This involved subtracting the power observed in the baseline condition from that in the learning condition, obtaining a pure learning effect characterized by relative changes in power. We categorized these power differences into three components: tagging responses tracking the high-probability location, tagging responses tracking other low-probability locations, and alpha oscillation separately. As depicted in Fig. 2B, as time progressed, tagging responses occurred in the intervals of 180 – 2238 ms and 2282 – 3192 ms for low-probability locations, while occurred in the interval of 1126 – 1862 ms for the high-probability location (cluster-based permutation test, *p* < .05; see Fig. S1 in Supplementary Information for the baseline condition), primarily falling within the pre-search time window (0 – 1250 ms). Notably, the difference between tagging responses towards high- and low-probability locations (referred to as the low-high difference, calculated by subtracting tagging responses towards those high-probability locations from that towards low-probability location) occurred in the intervals of 714 – 1442 ms and 1580 – 2252 ms (cluster-based permutation test, *p* < .05; see Fig. 2C). This indicates that participants can discern the high-probability location from other low-probability locations even before the commencement of the search, by showing a stronger frequency power in tagging response towards the high-probability location. Moreover, alpha activity was reduced in the learning condition in the intervals of 0 – 386 ms and 448 – 1812 ms relative to the placeholder onset (cluster-based permutation test, *p* < .05; see Fig. 2D). These results suggest proactive attentional mechanisms modulating high- and low-probability locations separately.

Additionally, this discrimination in tagging responses between high- and low-probability locations aligns with participants’ behavioral responses. Utilizing a median-split based on behavioral indices of learning (i.e., mean RTs for presenting a distractor in low-probability locations minus those in the high-probability location), we categorized participants into “superior learners” and “inferior learners” (n = 14 for each group). The results unveiled that the significant low-high difference persisted for superior learners in the interval of 246 – 2192 ms, while became weak and did not reach significance for inferior learners (see Fig. 2E left panel). However, the pre-search alpha activity existed for both superior learners (842 – 1598 ms) and inferior learners (720 – 1766 ms), with no significant difference detected between the two groups (cluster-based permutation test, *p* < .05; see Fig. 2E right panel). These findings underscore of the role of tagging responses in distinguishing high- and low-probability locations within learned suppression. Yet, the role of alpha oscillation remains ambiguous. We will return to this discussion later.

Since we consistently administered the baseline condition before the learning condition, to avoid the potential influences from learned suppression, one might question whether the observed results above were attributable to factors such as fatigue, practice effects, or any consequences associated with the fixed testing order. To address this concern, we partitioned the 10 blocks in the learning condition into the first 5 and second 5 blocks. If the observed effects were linked to the testing order, differences between the first 5 and second 5 blocks would be anticipated. However, no such distinctions were observed for alpha oscillation or the low-high difference (see Fig. 3). Notably, we also applied an inverted encoding model (IEM; Foster et al., 2016) to track the processing of target and distractor locations. Consistent with previous studies (e.g., van Moorselaar et al., 2020), our decoding results revealed that, through alpha activity, we could reliably track the target location but not the distractor location (see Fig. S2).

**Figure 3.**
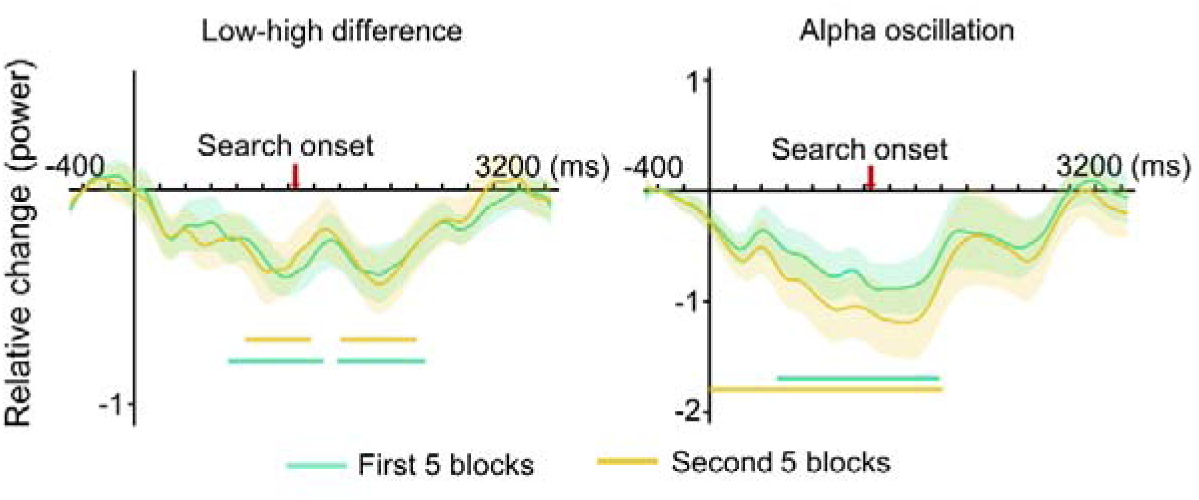
The low-high difference (left panel) and alpha oscillation (right panel) for first and second 5 blocks in the learning condition.

## Discussion

The present study aimed to investigate the neural mechanisms underlying the learned suppression of salient distractors using the frequency tagging technique (Ding et al., 2006; Malinowski et al., 2007; Morgan et al., 1996; Muller et al., 2003). Behaviorally, participants demonstrated proficient learned suppression, as evidenced by slower RTs for distractors presented in the high-probability location compared to low-probability locations. Importantly, as observed in tagging responses and alpha oscillation, the results unveiled the anticipatory neural mechanism underpinning learned suppression. This was evidenced by stronger tagging responses towards the high-probability location relative to low-probability locations, as well as a general reduction in alpha power before the search onset. Additionally, the ability to differentiate the high-probability location from other low-probability locations before the search onset primarily observed in participants with better learning performance.

It is generally agreed that learned suppression acts on the spatial priority map which ultimately determines attentional selection in an all or nothing manner (Duncan et al., 2023; Fecteau & Munoz, 2006; Ferrante et al., 2018; Sprague & Serences, 2013). This map represents a topographic space encoding the priority of individual locations by integrating signals from sensory input, current goals, and statistical learning (Theeuwes et al., 2022; Zelinsky & Bisley, 2015). According to this framework, if locations are likely to contain a target the location is up-regulated within the spatial priority map, whereas locations with a higher probability of containing distracting information are downregulated, resulting in the observed learned suppression.

Noted that, in the present study, we did not employ tagging for the search elements; instead, the placeholders (i.e., the stimulus background) were tagged continuously throughout the trial, with participants explicitly instructed to ignore them. Unlike traditional frequency tagging research, which typically involves continuously tracking attention at a specific location (Ding et al., 2006; Morgan et al., 1996), our experiment required participants to constantly shift their attention among four locations. Consequently, this setup led to a disconnect between the stimuli and their background. In other words, the background (i.e., tagging placeholders) served as a distraction for the search task. Previous studies have shown that enhancing attention to a stimulus causes the suppression of unattended (distracting) stimulus simultaneously (Andersen & Muller, 2010; Müller & Hübner, 2002), thereby leading to reduction in frequency power when tagging the unattended stimulus. This may explain the negative power observed for tagging frequencies in our study (Fig. 2A), as the tagging placeholders were disregarded throughout the trial.

Moreover, lower negative tagging responses to placeholders indicate that more attentional resources were allocated to suppressing placeholders (Andersen & Muller, 2010; Müller & Hübner, 2002). Since, at any moment in time, the total amount of attentional resources is limited, allocating fewer resources to processing search elements leaves more available for suppressing placeholders. Thus, lower negative tagging responses to placeholders reflect fewer attentional resources directed toward processing search elements, a reversed pattern compared to processing placeholders. In the learning condition, compared to the baseline condition, participants likely allocated less attention to the task, as they had already learned the high-probability distractor location, resulting in more attention to processing placeholders by showing lower tagging responses. This was further evident in behavioral findings, where overall mean RTs in the learning condition (1057 ms) were significantly faster than in the baseline condition (1116 ms), *p* < .001, suggesting that the learning condition was generally easier for participants, requiring less attentional focus. Accordingly, the lower tagging frequency power observed at low-probability locations relative to high-probability locations (Fig. 2C) indicates that participants allocated fewer attentional resources to processing stimuli at low-probability locations. Conversely, participants directed more attentional focus to stimuli presented at the high-probability location before the search onset. This anticipatory focus likely facilitated the suppression of the high-probability location immediately after search initiation.

This finding appears to contradict earlier studies suggesting that learned suppression operates proactively, without requiring prior attentional deployment to distractor locations (Gaspelin et al., 2015, 2017; Huang et al., 2022; Kong et al., 2020). For instance, Huang et al. (2022) demonstrated that in the context of statistical learning, reaction times to probes at high-probability distractor locations were already suppressed before display onset, consistent with proactive suppression. In contrast, more recent evidence challenges this view. Using a variant of the search-probe paradigm similar to that of Huang et al. (2022), Chang et al. (2023) demonstrated that the high-probability location was suppressed relative to all other locations. However, despite this suppression during search, they observed enhanced—rather than reduced—probe discrimination accuracy at this location in the probe task. This suggests that the high-probability location was initially attended before it was suppressed supporting a reactive suppression mechanism (e.g., Beck et al., 2018; Makovski, 2019; Moher & Egeth, 2012; Won et al., 2019; Chang et al., 2023; Chen et al., 2025).

Furthermore, Chen et al. (2025) employed the same task as we have used here while concurrently measuring micro-saccades, a well-established marker of covert attention. Consistent with the current and previous findings (Wang & Theeuwes, 2018a, 2018b, 2018c), they found that the high-probability distractor location was suppressed relative to all other locations. Yet, this learned suppression was accompanied by increased micro-saccade rates prior to stimulus onset, directed toward the high-probability location. This indicates that covert attention was initially allocated to the distractor location before suppression occurred, providing strong evidence for a reactive suppression mechanism. Taken together, these findings consistent with the present results suggest that covert attention is first allocated to distractor locations before suppression is engaged, reinforcing the view that suppression mechanisms may operate reactively rather than purely proactively.

Using EEG recordings, Wang et al. (2019) observed a relative increase in pre-search alpha power for electrodes contralateral to the high-probability location, providing direct neural evidence for proactive learned suppression as well. However, this observation contrasts with findings from other related studies (Ferrante et al., 2023; Noonan et al., 2016; van Moorselaar et al., 2020; van Moorselaar & Slagter, 2019). For instance, a recent magnetoencephalography (MEG) study showed that pre-stimulus neural excitability was reduced in the lateralized early visual cortex associated with the high-probability distractor locations, indicating the recruitment of proactive attentional mechanisms (Ferrante et al., 2023). However, Ferrante et al. did not observe alpha activity linked to learned suppression. A possible explanation for this discrepancy is that the learned suppression mechanism may require more complex processing to effectively suppress upcoming distractors in more demanding search tasks, which could involve alpha oscillation to support learned suppression. This may not apply when search is relatively easy. Indeed, previous research indicates that the modulation of alpha power, indicative of distractor suppression, is influenced by the perceptual load (Gutteling et al., 2022). Ferrante et al. (2023) adopted a rapid, invisible frequency tagging method in a search task that was comparatively less challenging, potentially accounting for the lack of alpha oscillation effects observed in their study. While the current study observed proactive alpha activity, its role may differ from that of tagging responses associated with learned suppression. This distinction highlights the need for further research to clarify the specific contributions of alpha oscillations and tagging responses in learned suppression mechanisms.

In sum, our findings provide direct neural evidence supporting the necessity of attending to distractor locations prior to suppression, indicating a reactive suppression mechanism.

## Notes

This research was supported by the Natural Science Foundation of Guangdong grant (2023A1515012789) to BW.

https://github.com/zhouyinfei2025/Shared_code

### Competing Interest Statement

The authors have declared no competing interest.

