## Supplementary information for "Pre-search attentional focus supports learned suppression in visual search"

**Time course of the baseline condition**

The primary aim of our study was to investigate the differences in tagging frequency power between high- and low-probability distractor locations. Since both conditions share the same baseline, we initially focused on comparing their time courses without explicitly including the baseline condition. However, in response to the reviewer’s insightful suggestion, we examined whether the baseline condition exhibits a similar time course to that of the high- and low-probability distractor locations. This analysis helps determine whether the observed temporal dynamics are specific to distractor location learning or if similar patterns emerge in a regular search task without statistical learning of distractor locations.

In Figure S1, we present the time-resolved power averaged across four flickering frequencies (2.4 Hz, 4.29 Hz, 5.45 Hz, 7.5 Hz) for the baseline condition, as well as the tagging responses corresponding to high- and low-probability distractor locations in the learning condition. Overall, the baseline, high-probability, and low-probability conditions exhibit a similar temporal trend, characterized by a general decrease in power over time. Specifically, all three conditions show significant power reductions compared to zero: the baseline condition exhibits a significant negative cluster from 438 to 900 ms, the high-probability location from 408 to 1394 ms, and the low-probability location from 426 to 2398 ms. However, despite this shared trend, significant differences exist among them. Direct comparisons reveal that both the high- and low-probability locations show significantly lower power than the baseline condition, with significant clusters observed from 1036 to 1886 ms for high-probability locations and from 428 to 2120 ms for low-probability locations.

These results indicate that while the overall decreasing trend is present across the baseline, high-probability, and low-probability conditions, the power reduction is more pronounced in the learning condition. This suggests that the temporal dynamics observed in the learning conditions are not merely a general trend in visual search but are likely influenced by statistical learning.

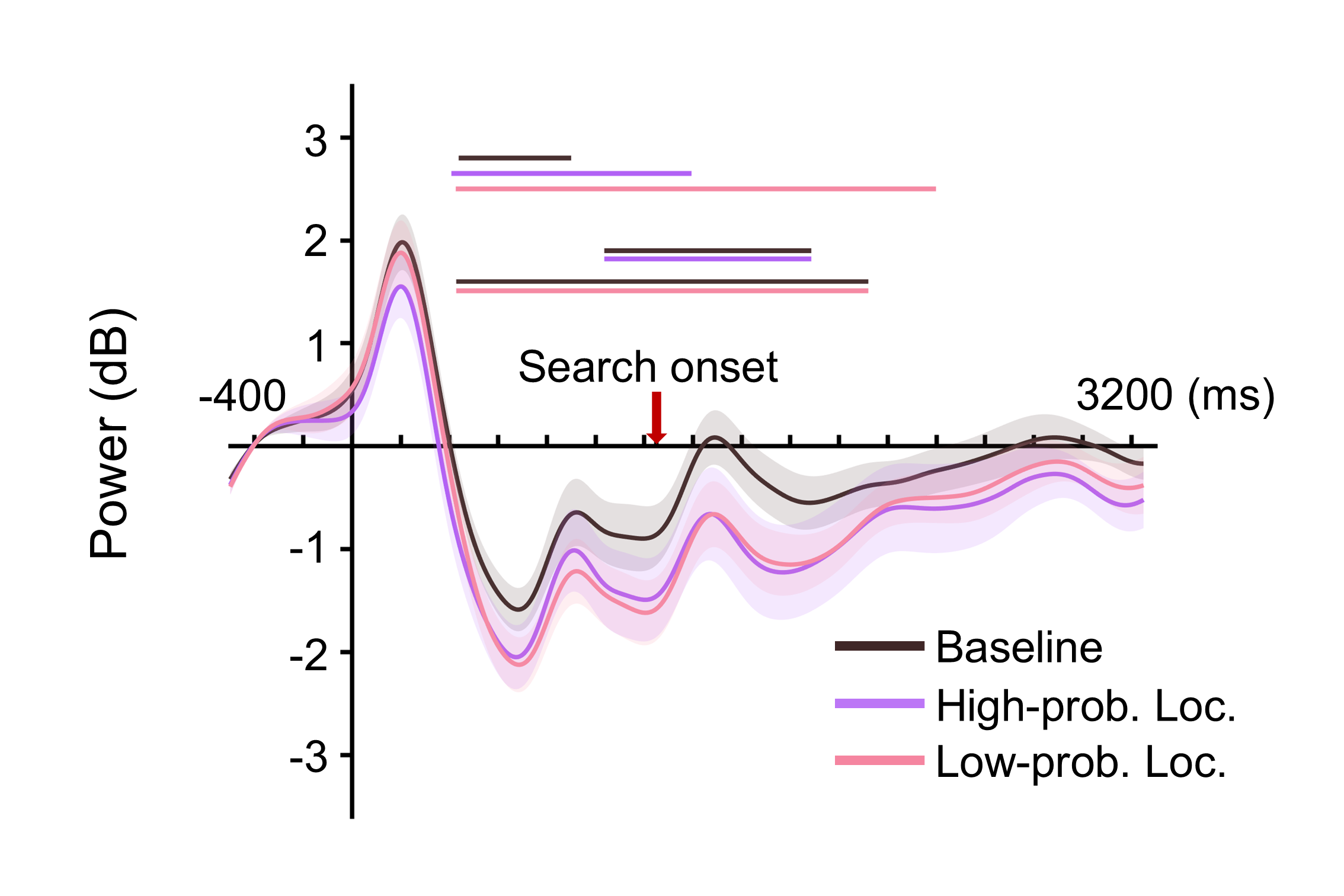

***Figure S1.*** Time-resolved power averaged across flickering frequencies (2.4Hz, 4.29Hz, 5.45Hz, 7.5Hz) in the baseline condition and for tagging responses corresponding to high- and low-probability distractor locations in the learning condition. Shadow areas represent ±1 SEM. Horizontal lines indicate significant clusters after cluster-based permutation test at *p* < .05.

**Inverted Encoding Model (IEM) analysis on alpha power**

To further investigate whether alpha oscillations encode spatial information about the high-probability distractor location, we conducted an inverted encoding model (IEM; Foster et al.,2016) analysis. This approach allows us to reconstruct spatially specific representations from EEG activity and determine whether alpha-band activity contains information about distractor locations.

This approach assumes that EEG activity at each electrode reflects a weighted sum of four spatially selective channels, each tuned to a specific location. Following standard IEM procedures, we modeled the response profile of each spatial channel using a raised cosine function, expressed as *R* = sin (0.5*θ*)^7^, where *θ* is the angular position (0°–359°), and *R* represents the channel response in arbitrary units. The model was circularly shifted to align each channel's peak response with a specific location. Data were partitioned into training (2/3 trials) and test (1/3 trials) sets using a cross-validation routine. Using training data, we computed a weight matrix W that mapped spatial channels to electrode activity: B = WC, where B is the EEG activity at the electrodes, and C is the predicted channel response. Using the estimated W, we reconstructed spatial channel responses from test data to obtain reconstructed CTFs.

To quantify spatial selectivity, we computed CTF slopes over time, with higher slopes indicating greater spatial specificity. If alpha-band activity encodes distractor locations, we would expect to see significant tuning for these locations. However, our results indicate that alpha activity reliably tracks the target position but does not encode the distractor location. Specifically, cluster-based permutation tests revealed significant clusters for target location tracking both in the baseline condition (2220-2482 ms) and the learning condition (1522-3054 ms), but not for distractor location tracking in either condition. This finding aligns with previous studies using spatial encoding models on EEG data (van Moorselaar et al., 2020), which demonstrated that expectations about distractor locations do not enhance spatial tuning to those locations. Instead, learned suppression mechanisms appear to reduce distractor-specific processing rather than encoding precise spatial information about distractors.

These results provide further support for the idea that alpha oscillations play a general role in learned suppression rather than in spatially specific distractor encoding. For completeness, we have included the results of this analysis in Figure S2, which illustrates the reconstructed representations based on alpha activity.

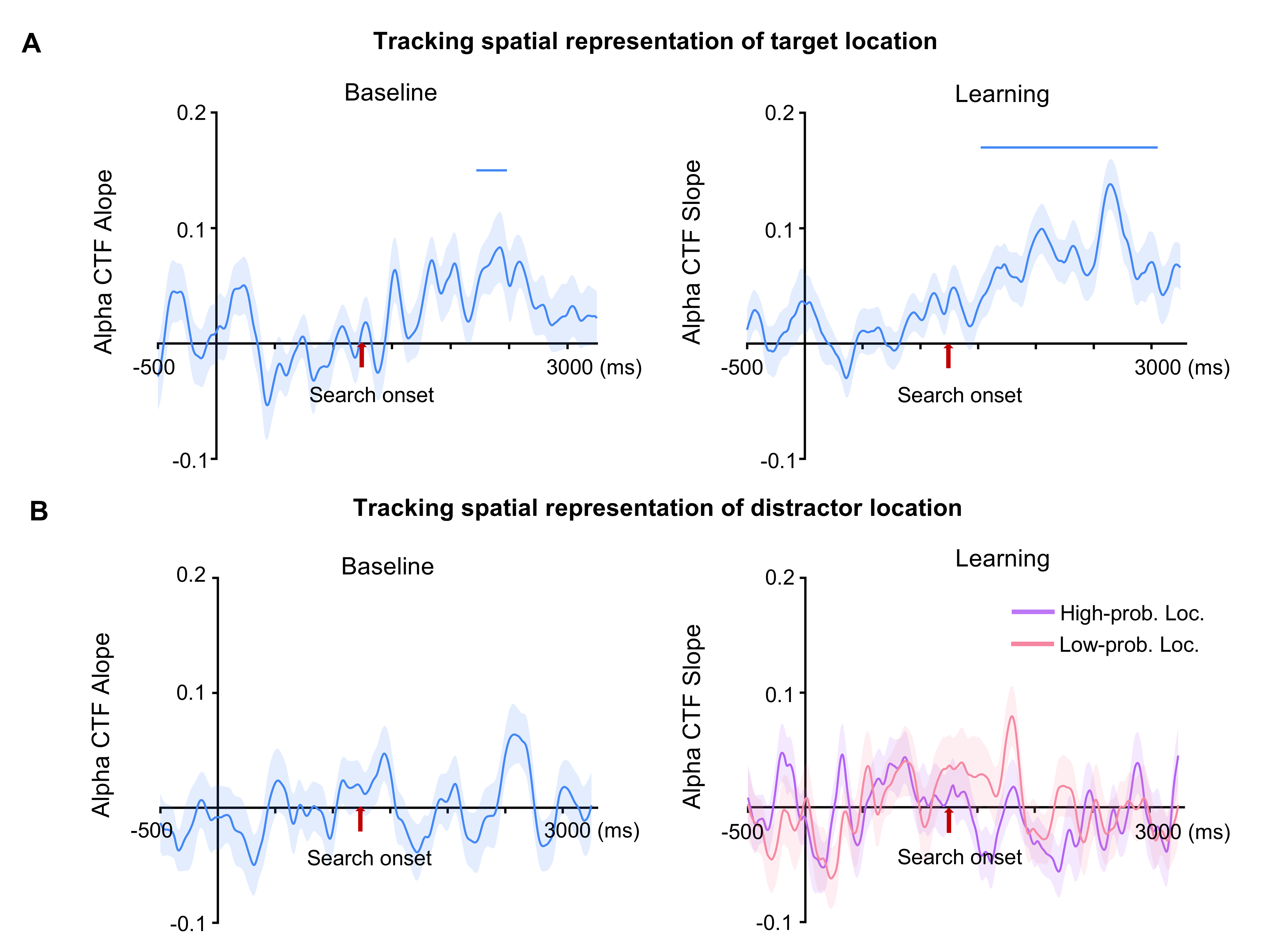

***Figure S2.*** Total power CTF slopes for tracking target (A) and distractor (B) locations. The CTF slope here quantifies the location specificity of the topographic distribution of alpha-band activity. The shaded areas represent ±1 SEM, and the horizontal lines indicate significant clusters where the CTF slopes significantly differ from zero after cluster-based correction (*p* < .05).

**Effect of cycle variation in time-frequency analysis**

To ensure that our time-frequency analysis results were not biased by the choice of wavelet cycles, we systematically varied the number of cycles between 1.5 and 2 times the corresponding frequencies. The power estimates obtained from different cycle settings were compared to assess consistency.

As shown in Figure S3, the power dynamics remained stable across different cycle parameters. The time courses and statistical comparisons exhibited negligible differences, confirming that our findings are robust to variations in wavelet cycle selection. These results support the validity of our chosen cycle range (logarithmically spaced between 3 and 12) and demonstrate that our key findings are not dependent on specific cycle settings (see Tables S1-S5 for detailed statistical values).

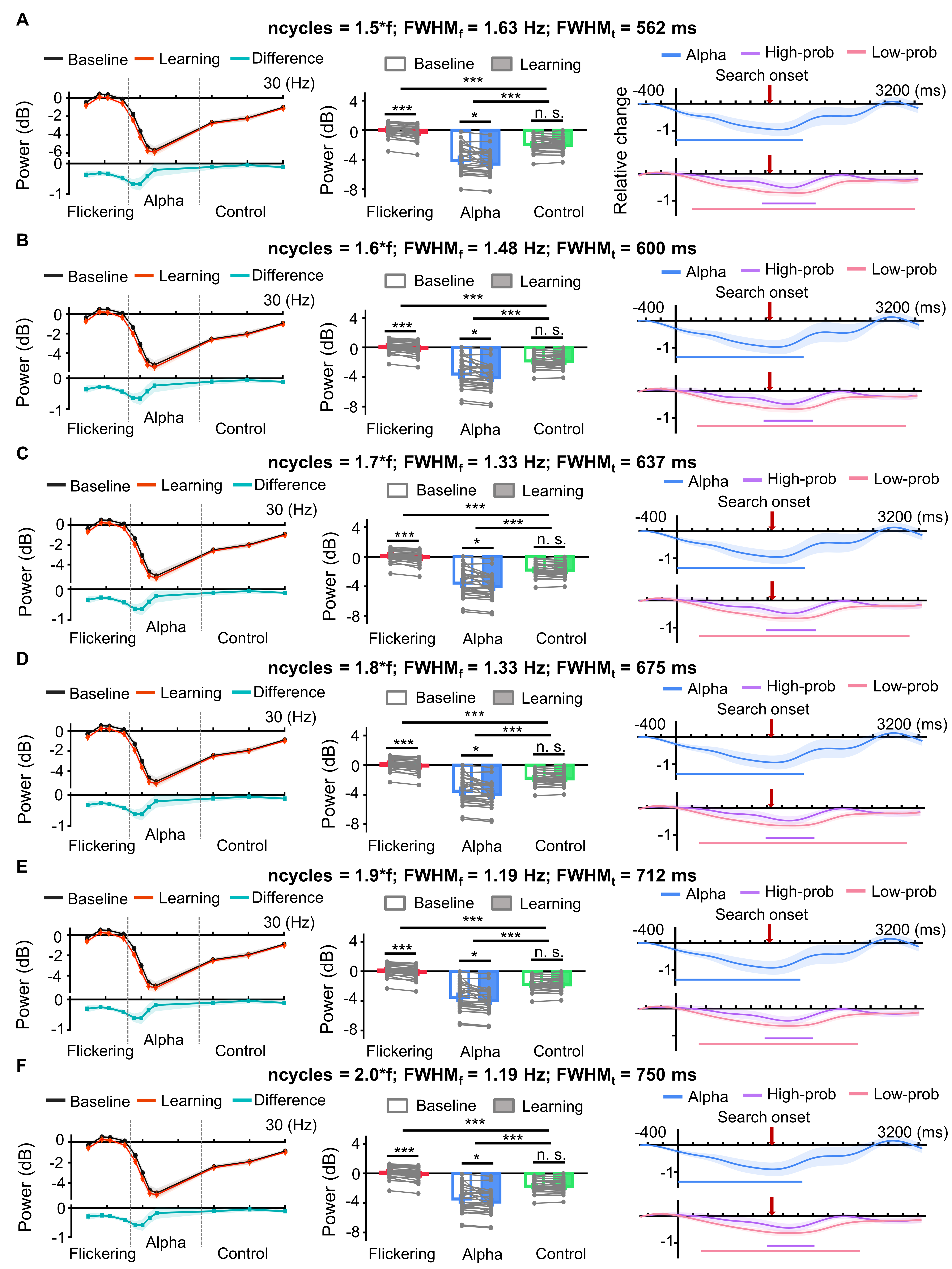

***Figure S3.*** Effects of varying the number of cycles in the time-frequency analysis. Panels A-F present the results of the time-frequency analysis performed with varying cycle numbers (1.5–2 times the corresponding frequencies), applied to the data depicted in Figures 2A and 2B of the main text.

**Tabel S1.** Summary of repeated-measures ANOVA results (Condition × Frequency type)

| Test condition | Effect | *F* | *p* | partial *η*^2^ |
| --- | --- | --- | --- | --- |
| A | Condition | 14.63 | < .001 | 0.35 |
|  | Frequency type | 97.28 | < .001 | 0.78 |
|  | Interaction | 3.68 | .032 | 0.12 |
| B | Condition | 16.22 | < .001 | 0.38 |
|  | Frequency type | 87.50 | < .001 | 0.76 |
|  | Interaction | 3.64 | .033 | 0.12 |
| C | Condition | 15.72 | < .001 | 0.37 |
|  | Frequency type | 88.43 | < .001 | 0.77 |
|  | Interaction | 3.54 | .036 | 0.12 |
| D | Condition | 15.06 | < .001 | 0.36 |
|  | Frequency type | 89.10 | < .001 | 0.77 |
|  | Interaction | 3.39 | .041 | 0.11 |
| E | Condition | 14.61 | < .001 | 0.35 |
|  | Frequency type | 90.17 | < .001 | 0.77 |
|  | Interaction | 3.26 | .046 | 0.11 |
| F | Condition | 14.06 | < .001 | 0.34 |
|  | Frequency type | 91.09 | < .001 | 0.77 |
|  | Interaction | 3.11 | .053 | / |

**Tabel S2.** Post-hoc comparisons of frequency type (baseline condition)

| Test condition | Comparison | *t* | *p* | *d* |
| --- | --- | --- | --- | --- |
| A | Flickering vs. Control | 8.40 | < .001 | 1.59 |
|  | Alpha vs. Control | 6.22 | < .001 | 1.18 |
| B | Flickering vs. Control | 8.91 | < .001 | 1.68 |
|  | Alpha vs. Control | 5.33 | < .001 | 1.01 |
| C | Flickering vs. Control | 8.90 | < .001 | 1.68 |
|  | Alpha vs. Control | 5.39 | < .001 | 1.02 |
| D | Flickering vs. Control | 8.86 | < .001 | 1.67 |
|  | Alpha vs. Control | 5.45 | < .001 | 1.03 |
| E | Flickering vs. Control | 8.83 | < .001 | 1.67 |
|  | Alpha vs. Control | 5.54 | < .001 | 1.05 |
| F | Flickering vs. Control | 8.79 | < .001 | 1.66 |
|  | Alpha vs. Control | 5.62 | < .001 | 1.06 |

**Tabel S3.** Post-hoc comparisons of frequency type (learning condition)

| Test condition | Comparison | *t* | *p* | *d* |
| --- | --- | --- | --- | --- |
| A | Flickering vs. Control | 7.41 | < .001 | 1.40 |
|  | Alpha vs. Control | 7.71 | < .001 | 1.46 |
| B | Flickering vs. Control | 7.93 | < .001 | 1.50 |
|  | Alpha vs. Control | 6.83 | < .001 | 1.29 |
| C | Flickering vs. Control | 7.94 | < .001 | 1.50 |
|  | Alpha vs. Control | 6.86 | < .001 | 1.30 |
| D | Flickering vs. Control | 7.94 | < .001 | 1.50 |
|  | Alpha vs. Control | 6.89 | < .001 | 1.30 |
| E | Flickering vs. Control | 7.95 | < .001 | 1.50 |
|  | Alpha vs. Control | 6.94 | < .001 | 1.31 |
| F | Flickering vs. Control | 7.94 | < .001 | 1.50 |
|  | Alpha vs. Control | 6.98 | < .001 | 1.32 |

**Tabel S4.** Post-hoc comparisons of condition differences

| Test condition | Comparison  (Learning vs. Baseline) | *t* | *p* | *d* |
| --- | --- | --- | --- | --- |
| A | Flickering | 5.83 | < .001 | 1.10 |
|  | Alpha | 2.60 | 0.015 | 0.49 |
|  | Control | 1.50 | 0.146 | / |
| B | Flickering | 5.50 | < .001 | 1.04 |
|  | Alpha | 2.73 | 0.011 | 0.52 |
|  | Control | 1.66 | 0.109 | / |
| C | Flickering | 5.49 | < .001 | 1.04 |
|  | Alpha | 2.69 | .012 | 0.51 |
|  | Control | 1.63 | .115 | / |
| D | Flickering | 5.47 | < .001 | 1.03 |
|  | Alpha | 2.62 | .014 | 0.50 |
|  | Control | 1.59 | .123 | / |
| E | Flickering | 5.49 | < .001 | 1.04 |
|  | Alpha | 2.57 | .016 | 0.48 |
|  | Control | 1.54 | .135 | / |
| F | Flickering | 5.45 | < .001 | 1.03 |
|  | Alpha | 2.52 | .018 | 0.46 |
|  | Control | 1.55 | .133 | / |

**Tabel S5.** Significant clusters identified by cluster-based permutation tests (*p* < .05)

| Test condition | Alpha activity | High-prob. Loc. | Low-prob. Loc. |
| --- | --- | --- | --- |
| A | [0 1706] | [1154 1872] | [216 3204] |
| B | [0 1708] | [1182 1848] | [290 3098] |
| C | [0 1696] | [1184 1844] | [294 3086] |
| D | [0 1684] | [1184 1836] | [298 3074] |
| E | [0 1670] | [1184 1830] | [304 2432] |
| F | [0 1656] | [1186 1822] | [310 2426] |
